# Development of a multi-locus typing scheme for an *Enterobacteriaceae* linear plasmid that mediates inter-species transfer of flagella

**DOI:** 10.1101/664508

**Authors:** James Robertson, Janet Lin, Amie Wren-Hedegus, Gitanjali Arya, Catherine Carrillo, John H.E. Nash

**Affiliations:** National Microbiology Laboratory, Public Health Agency of Canada, Guelph, Ontario, Canada; Ottawa Laboratory (Carling), Canadian Food Inspection Agency, Ottawa, Ontario, Canada; National Microbiology Laboratory, Public Health Agency of Canada, Toronto, Ontario, Canada

**Keywords:** plasmids, linear plasmids, serotyping, mobile genetic elements

## Abstract

Due to the public health importance of flagellar genes for typing, it is important to understand mechanisms that could alter their expression or presence. Phenotypic novelty in flagellar genes arise predominately through accumulation of mutations but horizontal transfer is known to occur. A linear plasmid termed pBSSB1 previously identified in *Salmonella* Typhi, was found to encode a flagellar operon that can mediate phase variation, which results in the rare z66 flagella phenotype. The identification and tracking of homologs of pBSSB1 is limited because it falls outside the normal replicon typing schemes for plasmids. Here we report the generation of nine new pBSSB1-family sequences using Illumina and Nanopore sequence data. Homologs of pBSSB1 were identified in 154 genomes representing 25 distinct serotypes from 67,758 *Salmonella* public genomes. Pangenome analysis of pBSSB1-family contigs was performed using Roary and we identified three core genes amenable to a minimal MLST scheme. Population structure analysis based on the newly developed MLST scheme identified three major lineages representing 35 sequence types, and the distribution of these sequence types was found to span multiple serovars across the globe. This MLST scheme has shown utility in tracking and subtyping pBSSB1-family plasmids and it has been incorporated into the plasmid MLST database under the name “pBSSB1-family”.

## Introduction

Serotyping is the current standard for classification of *Salmonella* isolates according to the reaction of antisera against the surface lipopolysaccharide layer (LPS) (O antigen) and flagellar (H antigens) (1–3). Based on the combination of antigens and biochemical characteristics an isolate is categorized into a serotype according to the White-Kauffman Le Minor (WKL) scheme (1–3). The *rfb* locus is important in determining the LPS layer phenotype but there is a complex genetic basis for O antigen phenotypes (4,5). The majority of *Salmonella* serovars possess two chromosomally encoded flagellar genes termed *fliC* and *fljB* that encode the H antigens. These flagellar proteins are alternately expressed as cells undergoing phase changes switch between transcription of the two genes (6). Phenotypic novelty in these important cellular components arise predominately through accumulation of mutations but horizontal gene transfer (HGT) is known to occur (4,7–9). An example of HGT affecting serologically important phenotypes is the plasmid mediated O antigen changes in the rare *Salmonella* serotypes Crossness and Borreze (10,11). Flagellar antigens have also been documented as being affected by HGT such as the case of *Salmonella* Typhi which normally expresses either the d or j flagella antigen (12,13) but a rare plasmid-borne variant expressing the z66 antigen exists (14). Baker et al. 2007b, discovered that the novel z66 flagellar gene was localized to a linear plasmid termed pBSSB1, which was able to mediate phase variation despite not being localized in the chromosome (15).

Whole genome sequencing (WGS) is revolutionizing the field of public health and it is replacing traditional serological testing as the primary diagnostic test for *Salmonella* and other pathogens (16). WGS provides an extraordinary level of discrimination of isolates, allows multiple tests to be run on the same data and provides a rich resource for the research community to answer novel questions which are not within the scope of traditional surveillance (17–19). However, the existing surveillance systems and historical data are dependent on serotype information and in order to maintain a connection to this important data, multiple tools have been developed for the purposes of predicting serotype based on sequence data (1,20). The *Salmonella in silico* Typing Resource (SISTR) identifies the genetic determinants for the O and H antigens from draft genome assemblies and uses 330 core gene to predict serotype with a high degree of accuracy (1,16). Presence of plasmid-encoded alleles of flagellar or O-antigen genes can confound WGS-based prediction of serotypes as these schemes currently do not account for the presence of multiple alleles of these genes..

Linear plasmids are extremely rare in *Enterobacteraceae* (15) and pBSSB1 is the only one known to occur in *Salmonella*. Typing of plasmids is traditionally based on replicon incompatibility where plasmids are grouped based on the ability to be stably maintained in a cell (21). The identification and tracking of this linear plasmid in bacterial populations is limited since pBSSB1 replicates through a different mechanism from the circular plasmids normally occurring in *Enterobacteraceae* and so falls outside the normal replicon typing schemes for plasmids currently in use. Multilocus sequence typing (MLST) is a technique for categorizing genetic diversity through assigning unique numeric identifiers for alleles of a set of genes which define the scheme (22). Traditional MLST schemes are based on a small subset of genes but the approach can be extended to any number of genes (1,23–25). MLST schemes have been developed for IncA/C, IncH, IncI and IncN replicon families, which facilitates the tracking of these plasmids through populations (26–29).

To date pBSSB1 had only been reported in *Salmonella* Typhi isolates from Indonesia presenting a z66 phenotype (14,15,30). Here we present a MLST typing scheme for the pBSSB1 plasmid backbone and information on the broad distribution of this plasmid in *Salmonella*. Based on phylogenetic analyses of the flagella and plasmid sequences, we have found evidence to support potential interspecies transfer of an intact flagellar operon from *Citrobacter* to *Salmonella*, which has implications for serology-based identification of *Salmonella*.

## Materials and Methods

### DNA preparation and sequencing

The OIE Reference Laboratory for Salmonellosis performed phenotypic serotyping according to accredited procedures. Genomic DNA was extracted using the Qiagen EZ1 robotic extraction system according to manufacturer’s instructions. DNA concentration was measured using the Invitrogen Qubit™ system, and quality of the DNA template was evaluated using the Agilent TapeStation™. Illumina MiSeq sequencing libraries were prepared using the NexteraXT kit according to the manufacturer’s protocol for 600-cycle sequencing. Nanopore sequencing was performed using the RAD002 or RBK004 rapid library preparation kit according to the manufacturer’s instructions on a R9.4 flow cell. Raw sequence data generated from this study was deposited into NCBI and the accession numbers are listed in Supplemental Table 1.

**Table 1.** Core genes from closed pBSSB1-family plasmid sequences were tested for selection using Tajima’s D statistic using MEGA 7 (m = number of sequences, n = total number of sites, S = Number of segregating sites, ps = S/n, Θ = ps/a1, π = nucleotide diversity, and D is the Tajima test statistic).

| Gene | Annotation | Average Length (bp) | Number of Alleles | m | S | ps | $\Theta$ | $\pi$ | D |
| --- | --- | --- | --- | --- | --- | --- | --- | --- | --- |
| group_13 | hypothetical protein | 410 | 6 | 12 | 47 | 0.11 | 0.04 | 0.05 | 0.91 |
| group_7 | hypothetical protein | 742 | 6 | 12 | 68 | 0.09 | 0.03 | 0.03 | 0.57 |
| *soj* | Chromosome-partitioning ATPase Soj | 626 | 6 | 12 | 126 | 0.2 | 0.07 | 0.08 | 1.29 |
| group_14 | hypothetical protein | 332 | 7 | 12 | 33 | 0.11 | 0.04 | 0.04 | 0.09 |
| *mqsA* | Antitoxin MqsA | 290 | 7 | 12 | 15 | 0.05 | 0.02 | 0.02 | -0.36 |
| group_1 | hypothetical protein | 695 | 8 | 12 | 85 | 0.13 | 0.04 | 0.04 | 0.17 |
| group_10 | hypothetical protein | 2333 | 8 | 12 | 362 | 0.16 | 0.05 | 0.06 | 0.7 |
| group_2 | hypothetical protein | 1121 | 8 | 12 | 143 | 0.13 | 0.04 | 0.05 | 0.5 |
| group_32 | hypothetical protein | 305 | 8 | 12 | 29 | 0.09 | 0.03 | 0.03 | 0.27 |
| group_33 | hypothetical protein | 344 | 8 | 12 | 29 | 0.09 | 0.03 | 0.03 | 0.27 |
| group_44 | hypothetical protein | 374 | 8 | 12 | 18 | 0.05 | 0.02 | 0.02 | 0.06 |
| group_8 | hypothetical protein | 254 | 8 | 12 | 32 | 0.13 | 0.04 | 0.04 | 0 |
| *higB-2* | Toxin HigB-2 | 353 | 8 | 12 | 14 | 0.04 | 0.01 | 0.01 | 0.06 |
| *traC* | DNA primase TraC | 1099 | 8 | 12 | 57 | 0.08 | 0.03 | 0.03 | 0.82 |

### Genome Assembly

Hybrid assembly using MiSeq and Nanopore reads was performed using Unicycler v. 0.4.5 with the default parameters (31). Each assembly was examined to confirm that every component was closed and circularized with the exception of the pBSSB1 plasmid. The terminal inverted repeats flanking pBSSB1-family plasmids were found to be difficult to assemble due to low sequencing coverage of the ends and the collapsing of repeats and assignment to either the 5’ or 3’ end of the plasmid (data not shown). This issue was not resolved by using Canu v. 1.8 (32), so the ends of the plasmids are likely incomplete. Each assembly was iteratively polished with Racon v 1.3.2 (https://github.com/isovic/racon) and Pilon v. 1.23 (https://github.com/broadinstitute/pilon) until no changes were made to the assembly. Unicycler with the default parameters was used to assemble publicly available MiSeq data for other isolates where long reads were unavailable in order to minimize variability due to differences in assembly procedure.

### *In silico* analysis of pBSSB1

Previously, we assembled 67,758 *Salmonella* genomes from the SRA (33) and each of these assemblies was checked for the presence of plasmids homologous to pBSSB1 (referred to hereafter as “pBSSB1-family plasmids”) using MOB-recon (34). The *Salmonella in silico* typing resource SISTR (1) was used to predict the serotype of each *Salmonella* assembly found to contain a pBSSB1 homolog. Serotypes for *E. coli* genomes were predicted using ECTyper v. 0.81 (https://github.com/phac-nml/ecoli_serotyping). MOB-recon reconstructed plasmids were annotated using Prokka 1.19 (35) and pangenome analyses were performed using Roary v. 3.12.0 with the identity threshold relaxed to 90% for core genes (36). A multiple sequence alignment for each gene was constructed using MAFFT v. 7.221 with the auto flag enabled (37). Tajima’s D statistic was calculated for each multiple sequence alignment using MEGA 7 with all three codon positions used (38). A maximum likelihood tree was generated for the concatenated multiple sequence alignments for each ST using MEGA 7 with the following parameters (100 bootstraps, Kimura 2-parameter model, gamma distributed rate, all coding positions). Population structure of the *Salmonell*a isolates was visualized using GrapeTree with the Enterobase cgMLST scheme (25,39). MLST allele calls were extracted using the MLST tool (https://github.com/tseemann/mlst) using the *S. enterica* or pBSSB1 schema based on the three genes *soj*, *higB* and *mqsA*.

### *In silico* flagellar gene analyses

Prokka 1.19 (35) was run on the sequences of pBSSB1-family plasmids which had been reconstructed using MOB-recon v. 1.4.8 (34) and genes annotated as “Flagellin” were selected for further analyses. Identical and truncated subsequences were identified using cd-hit-est (40) using an identity threshold of 1. The resulting unique set of sequences was subject to clustering in a second round with cd-hit-est using a threshold of 0.9 to identify any similar flagella alleles.

## Results

### Closed pBSSB1-family plasmid analysis

Long read sequencing using Nanopore was performed on nine *Salmonella* isolates found to contain a pBSSB1-family plasmid based on their Illumina sequence data. These newly closed plasmid genomes were analyzed along with three sequences from NCBI (NC_011422: *Salmonella* Typhi, CP026380: *Salmonella* Senftenberg, CP023444: *Klebsiella pneumoniae*). The accessions for all newly generated sequences are available in Supplemental Table 1. The closed pBSSB1-family plasmids ranged in size from 26kb to 33Kb with an average GC% of 36%. Pangenome analysis using Roary estimated a core genome of 14 genes (Table 1). Gene synteny was visualized for the closed plasmids using EasyFig with the following blast parameters (evalue >= 1e^−8^, length>=1500bp, identity >=75%) (41) (Fig. 1). Overall, there is a conserved central core region of the plasmid but the ends of the plasmids carry significantly different sequence content. Only six out of the 12 plasmids contained a flagella gene (Fig. 1). The plasmids from isolates SA20061017 and SA20130280 are nearly identical across their length. The sequence CP026380 clusters tightly with our newly generated sequences 11-5006 and GTA-FD-2016-MI-02533-1 to GTA-FD-2016-MI-02533-3.

**Figure 1.**
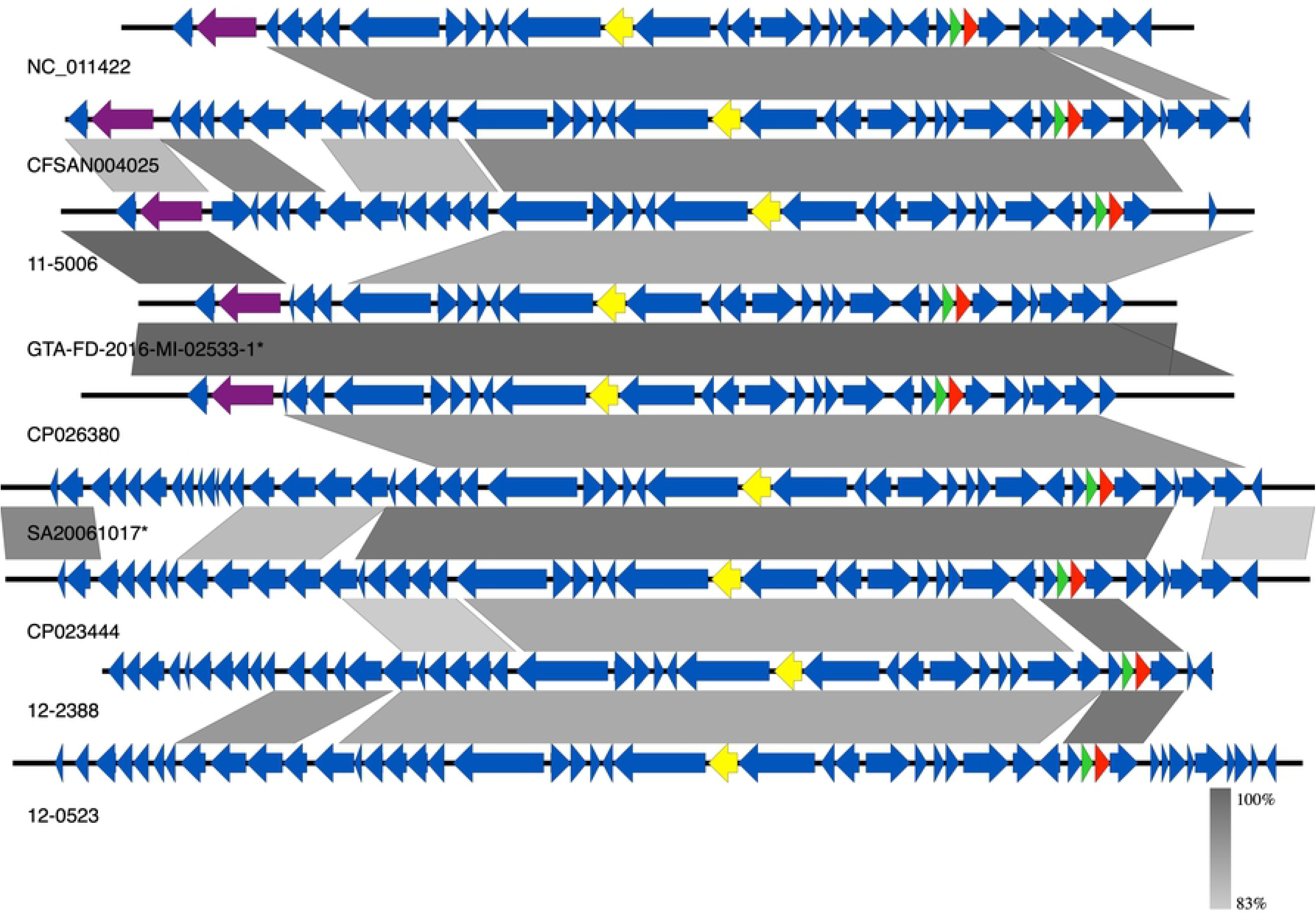
The sequence conservation for closed pBSSB1-family plasmids was visualized using EasyFig. Boxed arrows represent the position and transcriptional direction of ORFs. Shaded grey areas indicate conserved blocks with an evalue >= 1e-8. The locations of flagella genes are highlighted in purple. Genes associated selected for the three MLST scheme are highlighted in yellow (*soj*), green (*higB*), red (*mqsA*). Sequences with an asterisk indicate multiple samples with nearly identical sequences with a representative for that group: (SA20061017, SA20130280) and (GTA-FD-2016-MI-02533-1 to GTA-FD-2016-MI-02533-3).

### Development of a pBSSB1-family plasmid MLST scheme

In order to facilitate tracking of different lineages of the pBSSB1-family plasmid backbone, we developed a minimal MLST scheme based on its plasmid sequences. The distinct number of alleles for each of the core genes was determined and is listed in Table 1. Nine of the genes had 8 alleles with the remaining genes having either 6 or 7 alleles. Each of 14 core genes was tested for neutral evolution using Tajima’s D test in MEGA v. 7 (Table 1). None of the genes showed strong evidence for selection with *soj* showing the highest deviation from neutral with a Tajima’s D of 1.2 (Table 1). Since no significant selective pressure was observed for the core genes, all of them were considered viable MLST candidates. We identified three genes, which were good candidates for use as typing markers. We selected the sporulation inhibition homolog *soj*, along with the bacterial toxin/antitoxin (TA) genes *higB* and *mqsA.* The gene set resulted in 8 MLST profiles for the 12 closed plasmid sequences. Genes that contained multiple indels were excluded as candidates for MLST marker genes. The developed scheme has been deposited into pubMLST (https://pubmlst.org/plasmid/) under the name “pBSSB1-family” using the BIGSdb platform (42,43).

### Distribution of pBSSB1-family plasmids

A total of 154 *Salmonella* genomes out of the 67,758 SRA genomes were found to contain pBSSB1-family plasmids based on the results of MOB-recon. Each of these positive isolates was typed according to the *S. enterica* MLST scheme and then with the newly developed scheme for pBSSB1-family plasmids (Supplemental Table 2). A total of 35 pBSSB1-family sequence types were identified in the dataset with five sequence types accounting for 75% of the pBSSB1-family plasmids (Fig. 2). A minimum spanning tree based on the Enterobase cgMLST scheme was constructed using GrapeTree and overlaid with the pBSSB1-family sequence type to determine if the predominant sequence types were due to repeated samples from genetically similar members of a serovar (Fig. 3).

**Table 2.** Blast result summary from NCBI web-blast using a single representative per flagella sequence cluster.

| Allele | Representative | Length | Closest<br>NCBI Hit | Hit Species | Total<br>Score | Query<br>Coverage<br>(%) | E-value | Percent<br>Identity<br>(%) |
| --- | --- | --- | --- | --- | --- | --- | --- | --- |
| 1 | SRR3606556 | 1578 | CP012554 | *C. portucalensis* | 3337 | 100 | 0 | 99.37 |
| 2 | SRR3372244 | 1572 | CP012554 | *C. portucalensis* | 1803 | 100 | 0 | 78.76 |
| 3 | ERR1764822 | 1527 | NC_011422 | *S. Typhi* | 2809 | 100 | 0 | 100 |
| 4 | SRR3210535 | 1341 | CP037734 | *C. freundii* | 873 | 61 | 1e<sup>-150</sup> | 84.57 |

**Figure 2.**
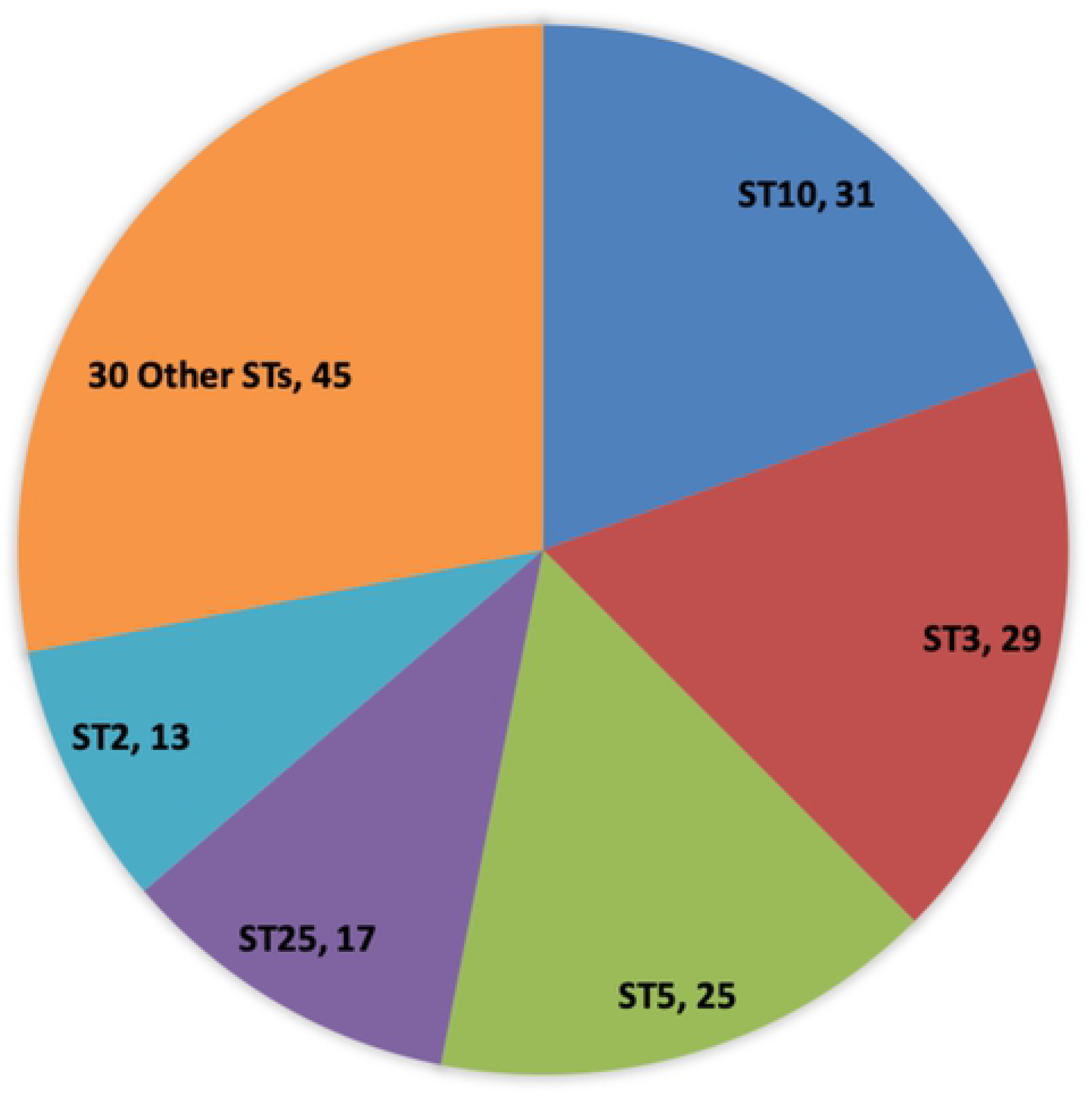
Pie chart indicating the MLST sequence type composition of identified pBSSB1-family STs in *Salmonella*. Counts of each sequence type are listed in each slice.

**Figure 3.**
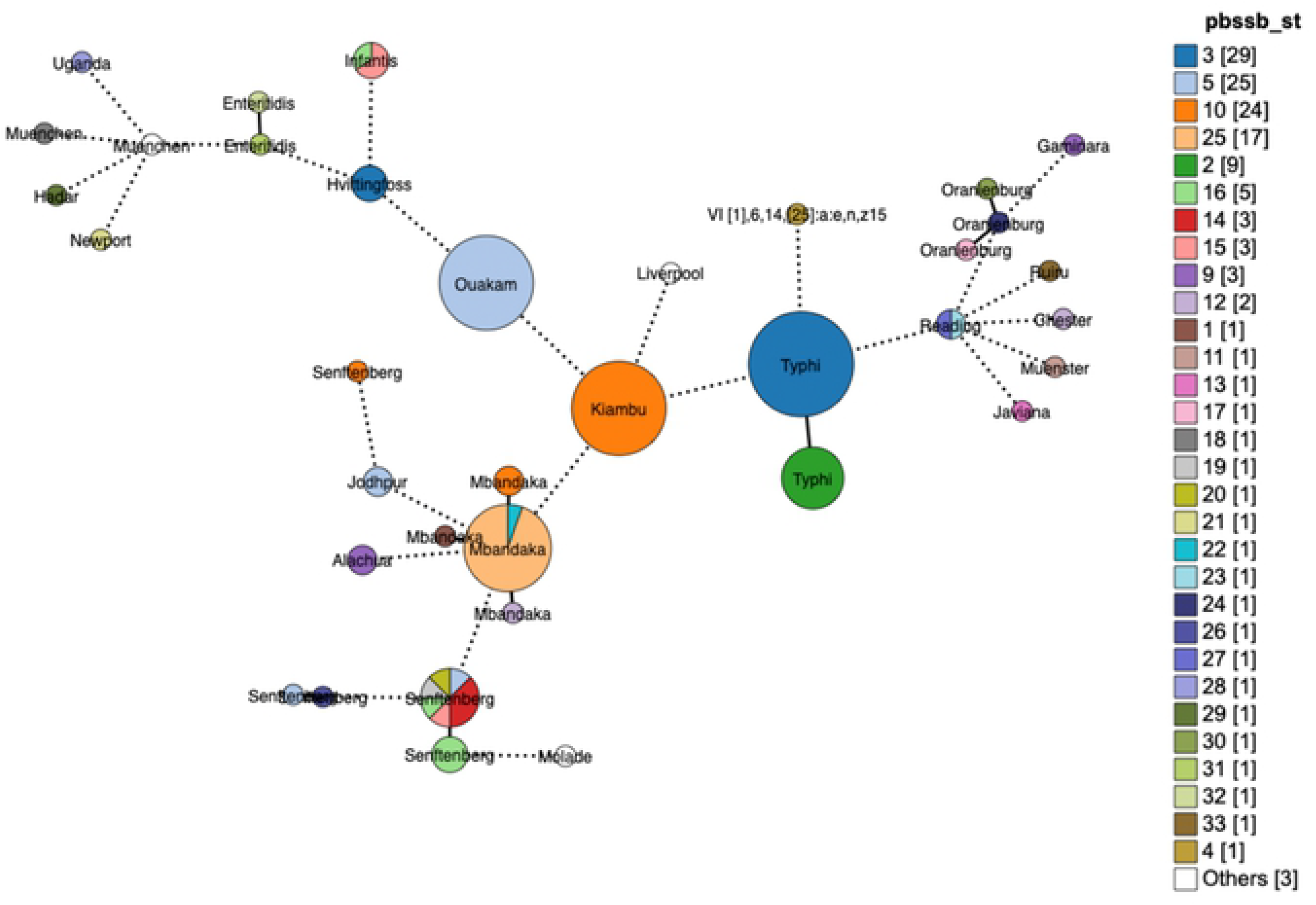
GrapeTree minimum-spanning tree based on the Enterobase cgMLST and colored based on the pBSSB1 sequence type present in the genome. Nodes differing by fewer than 50 alleles were collapsed together and branches longer than 500 alleles different were shortened and are indicated with a hashed line. Size of the nodes indicates the number of samples contained in them.

The pBSSB1-family MLST Sequence Type 10 (ST 10) primarily consists of serovar Kiambu isolates belonging to a single cluster (Fig 3), which is indicative of repeated sampling of closely related isolates. This pattern is consistent for the remaining isolates of ST 10 within different serotypes Mbandaka and Senftenberg (Fig. 3). A single cluster of Typhi isolates account for the majority of ST 3 isolates with a small cluster of Hvittingfoss accounting for the remaining three isolates (Fig. 3). A separate cluster of Typhi contains z66-positive ST 2, which indicates that not all pBSSB1 homologues in Typhi carry the z66 flagella (Fig. 3). A cluster of Ouakam contains the majority of ST 5, with isolates of Jodhpur and Senftenberg containing the others (Fig. 3). Infantis, Reading and Senftenberg are interesting cases because single clusters contain multiple pBSSB1-family sequence types (Fig. 3).

### Population structure of pBSSB1-family plasmids

A maximum likelihood tree based on the concatenated MLST gene sequences for each of the pBSSB1-family sequence types identified three major clades (Fig. 4). Both clades 1 and 2 contain significant sequence diversity, which is in contrast to clade 3 where the sequences form a tighter association. When the lineage information of pBSSB1-family plasmids is overlaid on the *Salmonella* population structure, there is evidence for both clonal expansion and horizontal transfer of lineages (Fig. 5). Each of the three different lineages are distributed across diverse serotypes (Fig. 5). The two clusters of Typhi contain either lineage 1 or 2 exclusively (Fig. 5). This is in contrast to Mbandaka, Senftenberg, Infantis and Reading where there are multi-lineage clusters occurring (Fig. 5). These results are consistent with repeated introductions of divergent plasmids into these serovars rather than spread and diversification of a single plasmid.

**Figure 4.**
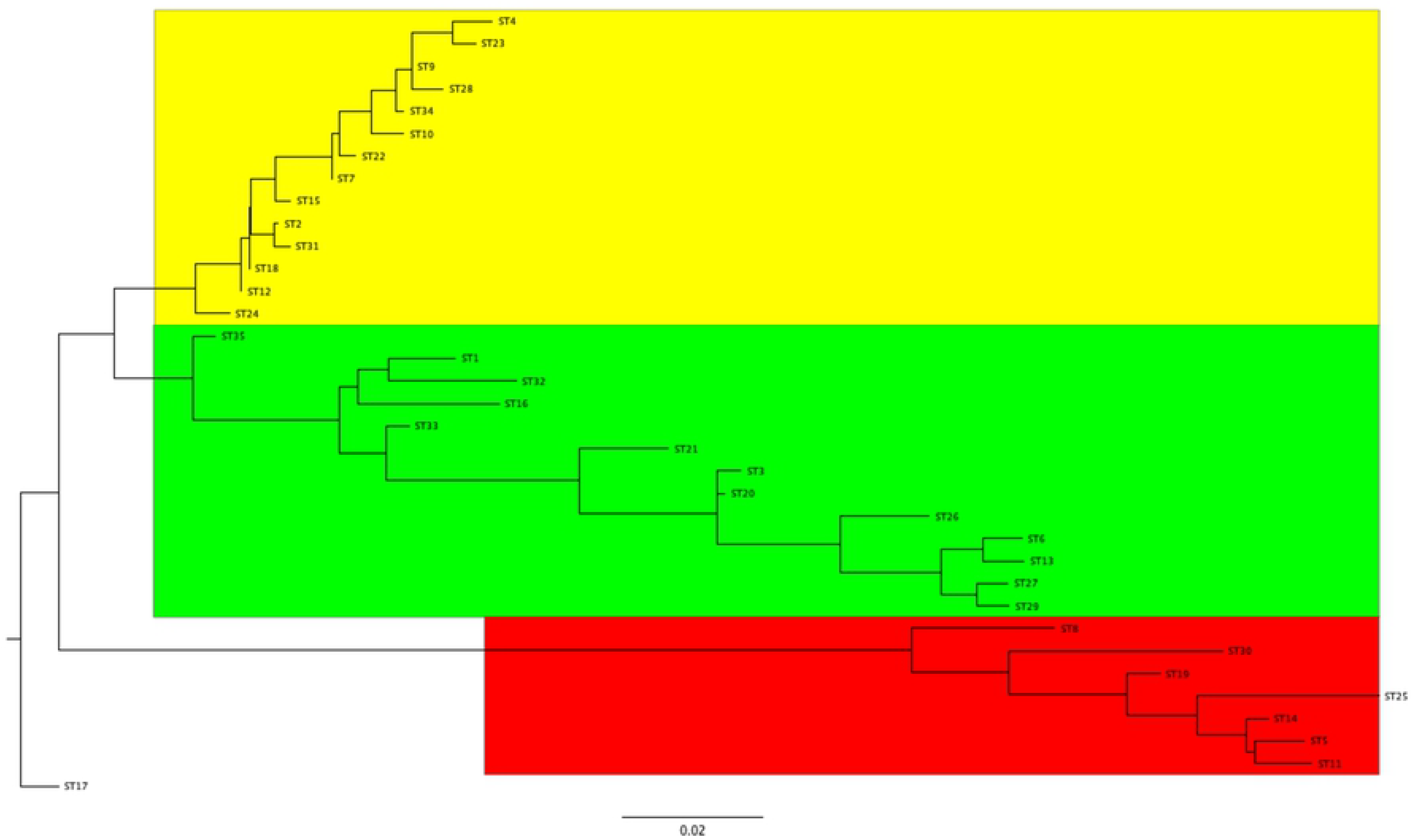
Maximum likelihood phylogenetic analysis of pBSSB1-family plasmids using concatenated sequences of the MLST genes *soj, mqsA, higB.* The sequence types have been divided into three major clades coloured in red (1), green (2) and yellow (3).

**Figure 5.**
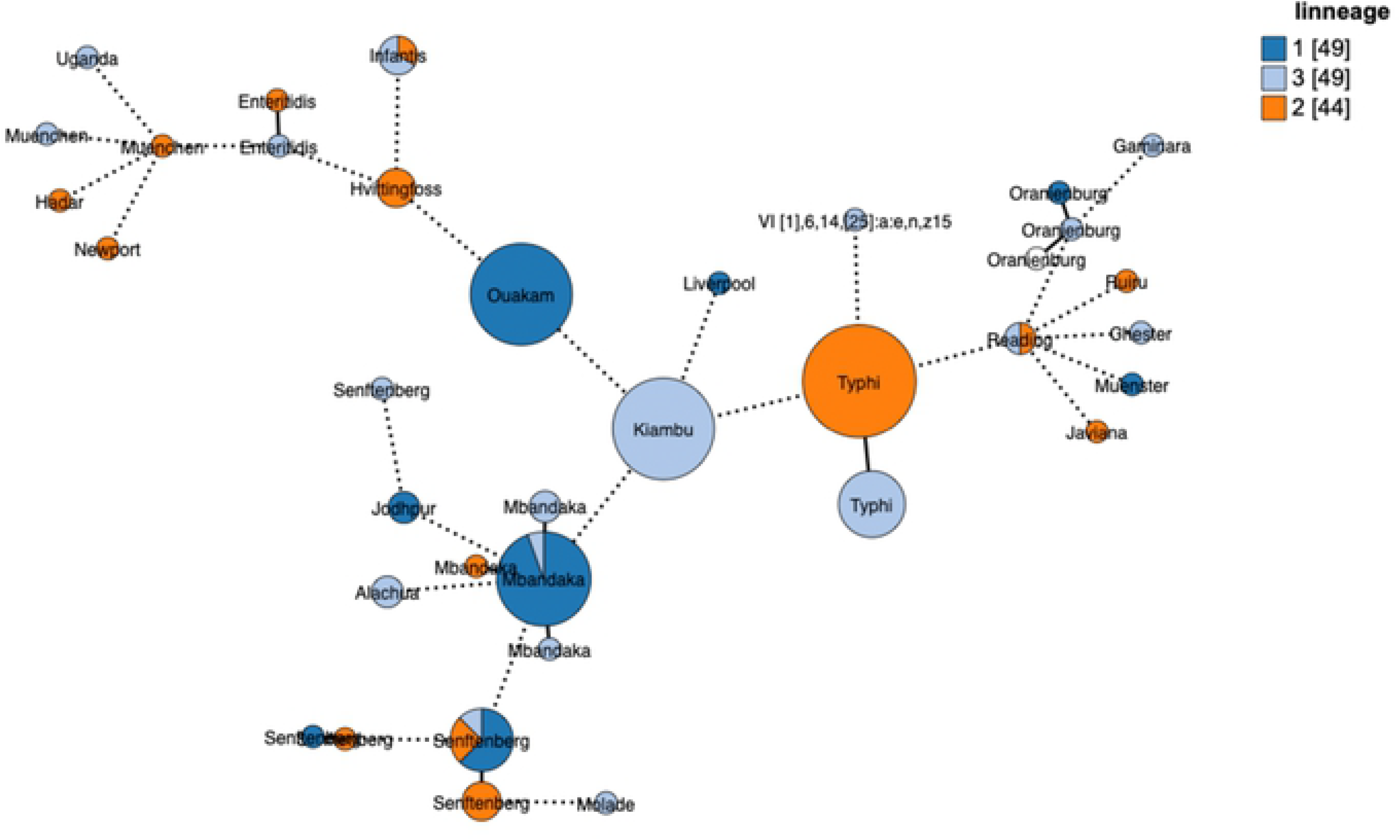
Grape Tree minimum-spanning tree based on the Enterobase cgMLST and coloured based on the pBSSB1-family lineages present in the genome. Nodes differing by fewer than 50 alleles were collapsed together and branches longer than 500 alleles different were shortened and are indicated with a hashed line. Size of the nodes indicates the number of samples contained in them.

### Plasmid mediated flagellar genes

Due to the presence of an intact *fliC* operon in some members of the pBSSB1-family, we examined the flagella sequences in detail to ascertain their similarity to other known *Enterobacteracea* flagella sequences. Flagellar genes were found in 104 of the 154 pBSSB1-family plasmids, which are distributed in 15 pBSSB1 STs and in all three lineages (Supplemental Table 2). There are total of 13 distinct flagella alleles including z66 from Typhi, which forms four clusters using cd-hit-est with a 0.9 threshold for identity. Web-based Blastn searches were performed using each of the allele sequences against the NCBI nucleotide database to identify possible sources of the flagellar genes (Table 2). Flagella cluster 1 and 2 both had their top hit as *C. portucalensis* (CP012554) but cluster 1 had much higher identity with 99.37% compared to 78.76% for cluster 2 (Table 2). Our samples 11-5006 and GTA-FD-2016-MI-02533-1 to GTA-FD-2016-MI-02533-3 belong to the flagella cluster 1 and our phenotypic serotyping results identified the z35 antigen but were unable to detect the normal g,[s],t flagella expression. This indicates that the genes encoding flagella on the identified pBSSB1-family plasmids are functional and these plasmid-encoded alleles are dominant relative to chromosomally-encoded flagellar genesand their presence masks the detection of the endogenous flagella. Sequences from cluster 1 share very little similarity with other z35 flagella in *Salmonella*, which is suggestive that there is cross-reactivity within the z35 antisera. Cluster 3 matched to the pBSSB1 plasmid NC_011422 from *Salmonella* Typhi and so represents the z66 flagella (Table 2). The fourth cluster matches with a chromosomal *C. freundii* flagella but overall had only 61% coverage and 84% identity (Table 2).

## Discussion

Given the importance of classification of *Salmonella* into serotypes, it is critical to characterize and understand the mechanisms, which generate novel antigenic combinations. The presence of variants of *Salmonella* Typhi containing a novel flagellar gene has been known since the 1980s (44), and in 2007 the linear plasmid pBSSB1 containing the z66 *fliC* was described (15). The plasmid pBSSB1 represents the only known vector for transferring an intact flagella operon in *Salmonella* and, based on the available data, it was only known to occur in Typhi isolates originating from some parts of Indonesia (15). This work represents the first description of pBSSB1 in diverse serovars and geographic locations. Analysis of 67,758 publicly available genomes from a previous study (33) shows that the plasmid is in fact globally distributed and present in a variety of serotypes (Fig. 2). The wide distribution of pBSSB1-family in a variety of serotypes and species indicates that this plasmid backbone could contribute to the generation of novel flagellar phenotypes through inter-species transfer. The transfer of this plasmid is known to be dominantly expressed over the endogenous *fliC*, which can result in incomplete typing of isolates by phenotypic methods (15). This is of concern to public health since serotype information is a critical piece of outbreak detection and response.

The circulating pBSSB1-family plasmids identified in this study represent diverse lineages rather than clonal spread of a single plasmid backbone (Fig. 2). The analysis using GrapeTree based on the Enterobase (25) cgMLST scheme overlaid with pBSSB1-family ST information, highlights that there has been repeated sampling of closely related isolates within serotypes (Fig. 3). Senftenberg is notable since within cgMLST clusters there exist multiple pBSSB1-family sequence types (Fig. 3). These results support the hypotheses that there were multiple independent acquisitions of the plasmid within this serotype. Estimates of the frequency of pBSSB1 homologues in *Salmonella* as a whole based on the SRA data should be undertaken with caution since the dataset is heavily biased towards repeated sampling of outbreaks and human clinical cases. However, given that pBSSB1 homologues were found in less than 0.3% of samples it is suggestive that it is not common within *Salmonella* of clinical relevance.

## Conclusion

This is the first documentation of plasmids similar to pBSSB1 outside of Indonesian *Salmonella* Typhi and provides evidence for global distribution. These results are of consequence to public health since serological classification of *Salmonella* is still the global standard and plasmids belonging to the pBSSB1-family can be vectors that can alter the flagellar phenotype of an isolate. These classification issues will still be present even after the public health reference laboratory community switches to WGS since serotype information remains critically important for investigations and reporting. The development of a pBSSB1-family MLST will aid in the tracking of these plasmids through different bacterial populations.

## Abbreviations

cgMLST: core gene multi-locus sequence typing
MLST: multi-locus sequence typing
ST: sequence type
WGS: whole genome sequencing
WKL: White-Kauffman Le Minor serotyping scheme

## Acknowledgments

We thank our colleagues within the National Microbiology Laboratory’s Reference Services Laboratory and the OIE Salmonella Reference Laboratory within the Division of Enteric Diseases for their assistance with phenotypic testing of the isolates. In addition, we would like to thank Paul Manninger for preforming WGS of some of the samples, Andrew Low for bioinformatics support, as well as Adam Koziol and Moe Elmufti for their comments and critiques during the review process. We also would like to thank the Food and Drug Administration, Center For Food Safety And Applied Nutrition (CFSAN) for providing the isolate of CFSAN004025. Finally, we would like to thank Marc Stevens and Dr. Roger Stephan from Institute of Food Safety, University of Zurich who provided the raw PacBio data for CP026380.

